# Methods in Description and Validation of Local Metagenetic Microbial Communities

**DOI:** 10.1101/198614

**Authors:** David Molik, Michael E. Pfrender, Scott Emrich

## Abstract

1. We propose MinHash (as implemented by MASH) and NMF as alternative methods to estimate similarity between metagenetic samples. We further describe these results with cluster analysis and correlations with independent ecological metadata.

2. Using sample to sample similarities based on MinHash similarities we use hierarchal clustering to generate clusters, simultaneously we generate groups based on NMF, and we compare groups generated from the MinHash similarity derived clusters and from NMF to those determined by the environment, looking to Silhouette Width for an assessment of the quality of the cluster.

3. We analyze existing data from the Atacama Desert to determine the relationship between ecological factors and group membership, and using the generated groups from MASH and NMF we run an ANOVA to uncover links between metagenetic samples and known environmental variables such as pH and Soil Conductivity.

## Introduction

How microbial communities, and in some broader context local communities, are determined, described, and validated is a matter of some debate (Holyoak et al. 2005). Principal Components Analysis (PCA) is the most common computational approach used to asses patterns of community. A typical metagenetic experimental design being the sequencing of gene regions that have gone through PCR from aquatic or soil samples, sequences would then be made OTUs, and PCA applied. PCA is biased towards components that have the most variance (Parsons et al. 2009). Communities also can be delineated from another by inferred differences in the identity and abundance of species detected within one or more samples (Rusch et al. 2007; Seshadri et al. 2007). Here we present two such alternative computational methods: MinHash (Broder, 1997) sketching and Non-Negative Matrix Factorization (NMF) (Seung & Lee 1999). NMF can be paired with k-means to estimate the number of groups (i.e., potential local communities) present and has the benefit of determining the most important feature driving inferred relationships. MinHash sketches can be used to quickly estimate similarities between whole samples in an alignment-free approach, i.e., OTUs do not need to be generated first. While MinHash and NMF are used here to cluster metagenetic samples based on inferred relationships, note that NMF focuses on what is distinct (in a cluster) while a MinHash implementation (Ondov et al. 2016) is combined with hierarchical methods to infer clusters based on pairwise similarities.

Because detected abundances of a species in metagenetic samples may not correlate with its actual abundance in a broader area, drawing boundaries between actual local communities using any computational approach can be difficult. As a result, prior work in community analysis has often relied on metadata such as physical barriers and environmental measurements to refine the structure of estimated local communities based on the species observed (Holyoak et al. 2005). We propose similar metadata-driven analysis using Silhouette plots for cluster assessment, which is a computational measure of how close each point in a cluster is to other clusters. Finally, we use ANOVA statistical tests to determine association of known environmental factors to the inferred clusters using our new and existing approaches. The advantage of these novel, data-driven (unsupervised) approaches for defining communities is that it allows us to artificially induce computational cutoffs, and, as a result, no prior knowledge/metadata are required to infer associations. Because environmental characteristics can change the viability of a microbial species occupying that area (Hultman et al. 2015; Gibbons & Gilbert 2015), subsequent comparisons of groupings to independent environmental variables provides a biologically motivated assessment of whether these computationally generated results uncover local communities.

To assess our new approachm we have chosen Atacama desert microbial community because of the data’s wide geographic range and inclusion of environmental variables. Log-likelihood statistical analysis of an indicates that among these *de novo* methods applied to Atacama data, hierarchical clustering using MinHash similarities has more explicative power than NMF on OTU abundance (see supplemental). In the previously reported analysis of this Atacama desert dataset samples taken from the same sampling location (North/Central/South) were more similar according to alpha diversity (Crits-Christoph et al. 2013); however, we show that other environmental variables can have a statistically higher correlation than sampling location, and specifically that pH, air relative humidity (RH) and soil conductivity best explain observed local communities derived computationally. Combined, these results indicate data-driven methods can be directly used to estimate community structure from NGS data.

## Methods

To define clusters we introduce MinHash (Ondov et al. 2016) based similarity for determining local community structure, which is essentially an approximation of the Jaccard similarity based on shared species within samples (see Rusch et al. 2007 and Ondov et al. 2016 for details). We also apply Non-Negative Matrix Factorization (NMF) (Gaujoux & Seoighe, 2010); (Seung & Lee 1999); (Paatero & Tapper 1994) using the nsNMF algorithm (Pascual-Montano et al. 2006) to determine non-shared species based on OTU abundances.

NMF—or Non-Negative Matrix Factorization—is method by which to split a matrix into a component based on the factors that are most important in making that split. For example, for RNA-seq expression analysis, suppose there are ‘k’ known clusters. NMF will break a provided expression matrix (genes by cells or cell tissues) into *k* total clusters while also producing the most important genes for doing so (Yu-Jui, 2017). When applied to observed OTU abundances, NMF will ideally return the most important OTUs to generate a fixed number of clusters. The power in this method is that different factors may be indicators for each cluster, instead of just the presence or absence of a particular observed species. NMF becomes particularly powerful when paired with k-means (Hartigan & Wong, 1979); (Forgey, 1965), which is a clustering method that can be used to measure how many clusters exist (aka, the ‘fit’). Determining factors in NMF can be done with Non-Negative coefficients while PCA has orthogonal vectors with positive and negative cofficients and since NMF combines factor discovery with iterative determination of the total number of clusters, NMF can be a more descriptive alternative to simple PCA-based visualization.

MASH, which is based on MinHash sketching (Broder, 1997), is an alignment-free method by which to estimate the distance between two sequences or sets of sequences. Using this computational method, a set of samples can be sequenced and then quickly compared to estimate how similar they are. The resulting pairwise similarity matrix can then be clustered hierarchically and visualized in the form of dendrograms and/or heatmaps. MASH can be run on raw samples at the cost of potentially higher inferred distances. Example hierarchical clustering algorithms are Diana (Struyf et al. 1997); (Kaufman & Rousseeuw, 2009) and McQuitty-WPGMA (McQuitty, 1966).

Using Silhouette Widths (Rousseeuw, 1987); (Handl et al. 2005) and the clustering information derived from NMF we can further describe structure within a cluster. Specifically, Silhouette Width highlights the ‘belongingness’ of each data point within a cluster; higher averages indicate cluster points are more tightly correlated with each other. Silhouettes are a tool to show how much overlap there is between clusters, or how consistent or distinct they are, similar to looking for distinct clusters in PCA plots.

Finally, we hypothesize that the local assortment of species is largely determined by the environment in which they live. If so, a change in environment and a corresponding change in observed species should, for the most part, correlate and this correspondence can be tested using both ANOVA and a mantel test under the right conditions (DeLong, 2013). We also realize that environment itself can correlate with distance, i.e., in the northern hemisphere, northern samples have fewer growing degree days than southern samples. For this reason isolation-by-distance (IBD) could also manifest as distinct clusters using our computational alternatives just as they would in a traditional PCA analysis.

## Results

Sample clustering based on OTUs was performed using Non-negative matrix factorization (NMF), which determines OTUs that are most informative using linear algebra-based techniques (Ondov et al. 2016; Seung & Lee 1999; Paatero & Tapper 1994; Yu-Jui, 2016). Sample to sample distances were determined based on minhash sketches, which estimate the Jaccard similarity of two samples based on shared subsequences (k-mers). We also determined the OTUs present in these Atacama samples using mothur (see Methods). Given our focus on unsupervised analysis, we processed the mash-based sample distances with multiple clustering methods: K-means (Hartigan & Wong, 1979; Forgey, 1965), hierarchical (Everitt, 1974; Hartigan, 1975), Agglomerative and Divisive (Kaufman & Rousseeuw, 2009).

**Figure 1.**
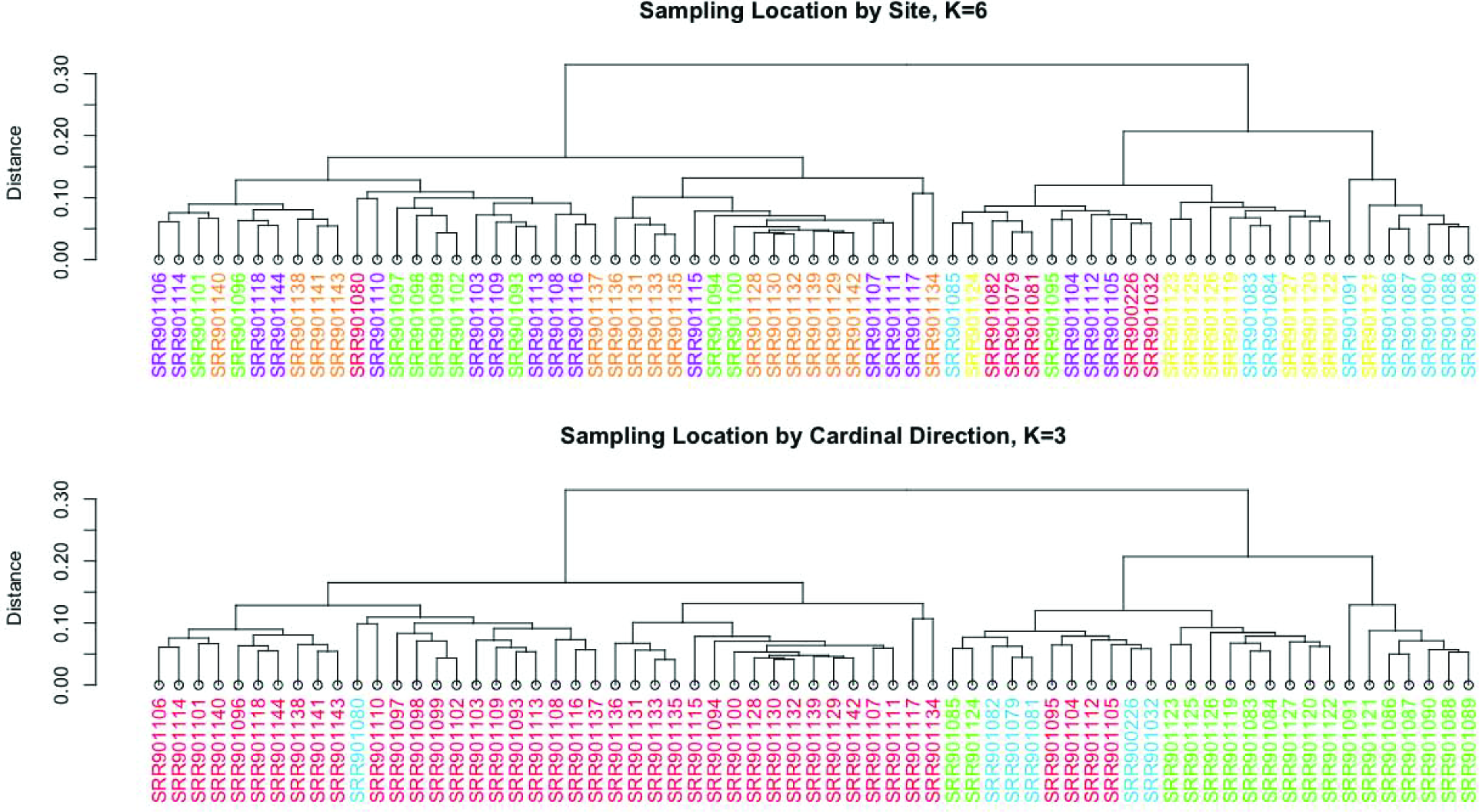
*Sample clusterings of Crits-Christoph et al. (2013) data using two measures of distance: site location (top) and cardinal direction (bottom). Dendrograms were generated with McQuitty Algorithm and are colored by sampling location (top, 6 total).*

**Figure 2.**
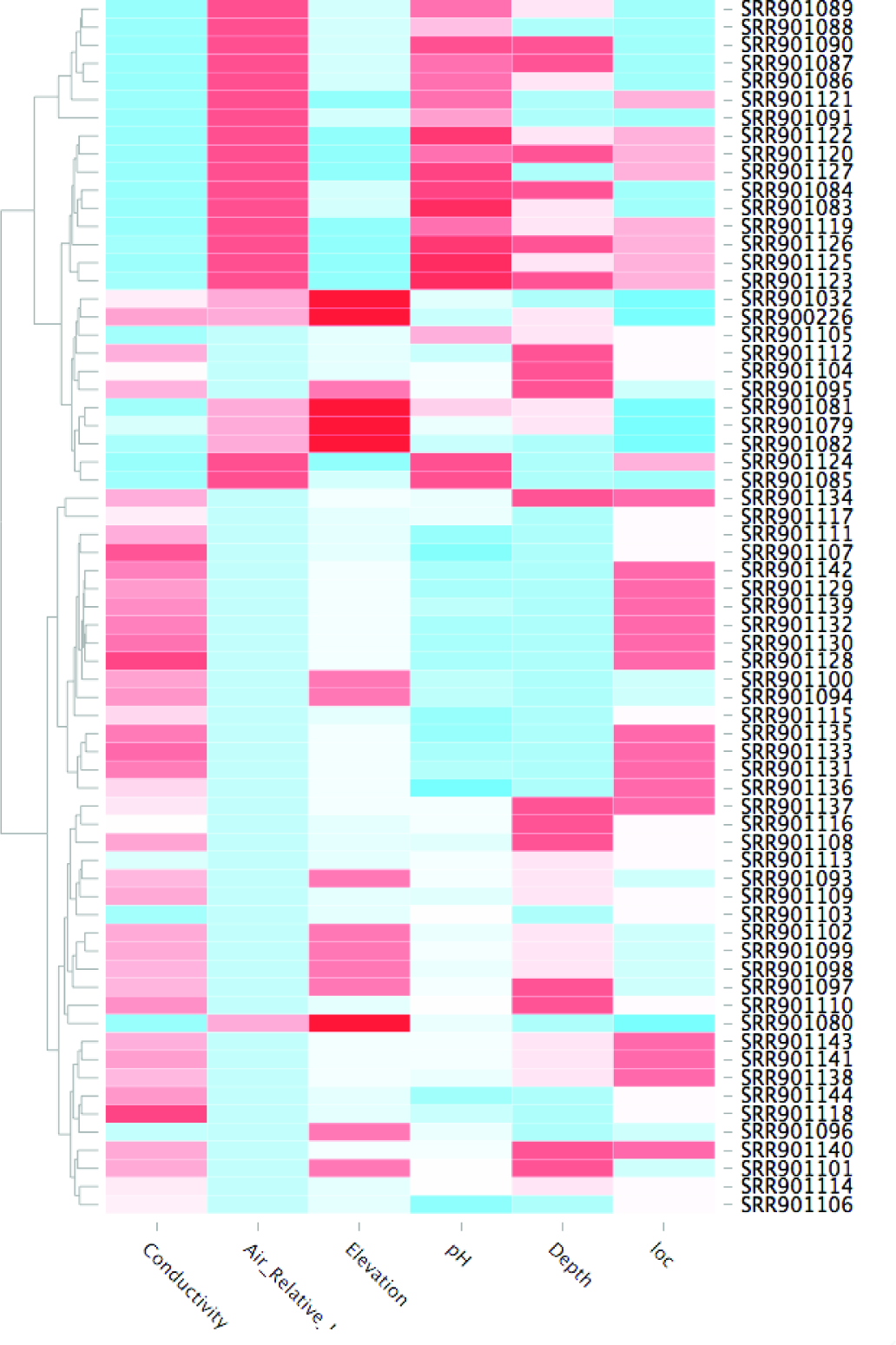
*Heatmap of Various Environmental Variables, scaled in color from low (blue) to high (red), colo scale per column. The left hand side is determined by McQuitty clustering algorithm on sample to sample similarities, as in Figure 1.*

Although prior work had shown that alpha diversity relationships among Atacama desert samples were driven by geographic location (Crits-Christoph et al. 2013), our preliminary analysis suggested sample to sample similarities based on mash and NMF were better explained by pH, Relative Air Humidity, and Conductivity as well as the previously reported location variable. Note that this “cluster first” computationally focused approach is a departure from previous techniques that draw local communities using external metadata to overcome species dispersion, although the species’ relationships are often defined by interrelated sequence clusters (OTUs).

**Figure 3.**
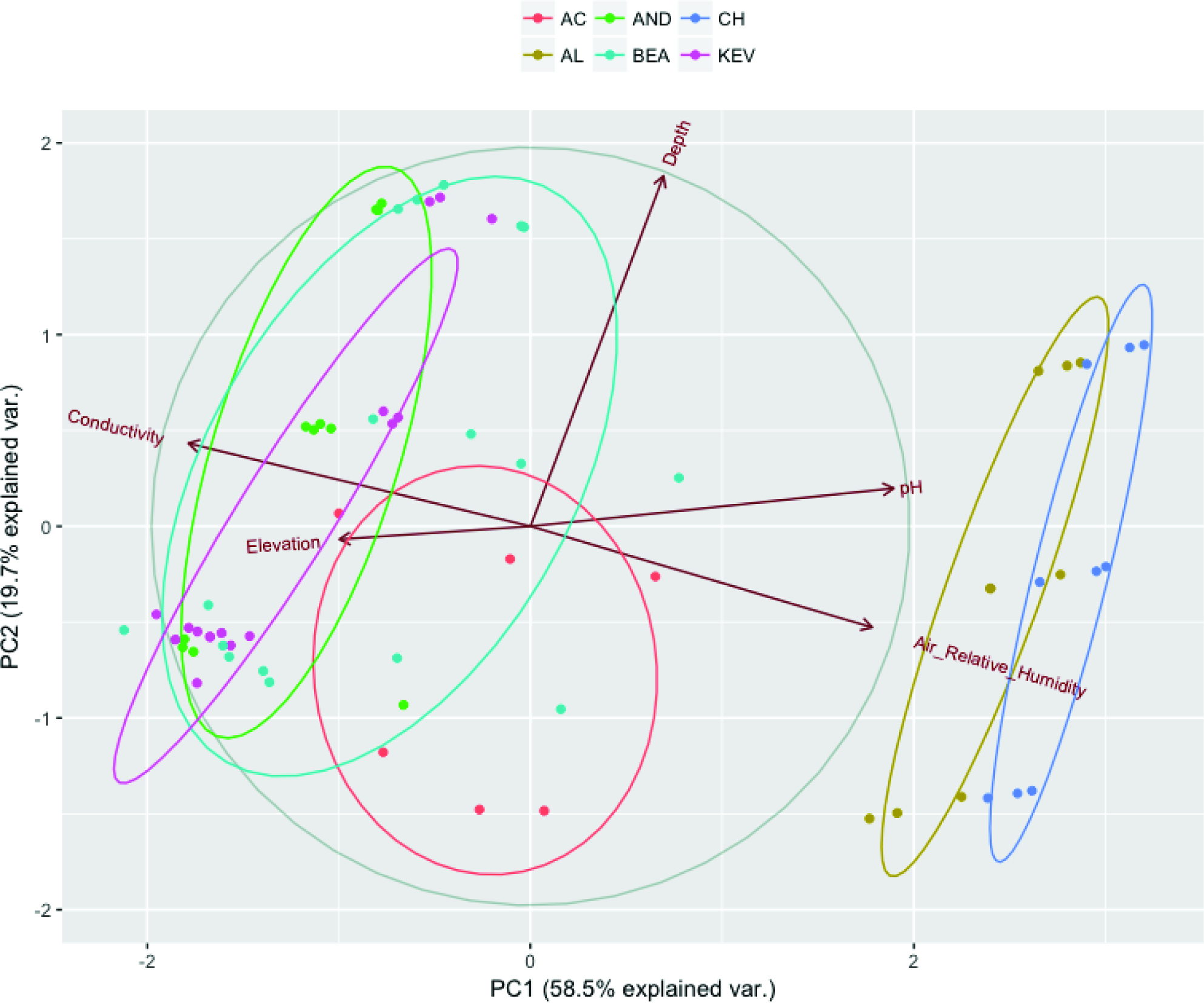
*PCA*(Hartigan & Wong, M 1979) *of environmental variables, colored according to sampling site*.

To be consistent with current practice, we first applied Principal Component Analysis (PCA) using the samples’ environmental variables (Air Humidity, Depth, Elevation, Soil Conductivity, and PH) to assess whether there is a ecological basis for observed clusters (Figure 2). We also used Average Silhouette Width (Rousseeuw, 1987), which provides a measure of how dense clusters are, with denser clusters being preferred. Average Silhouette can be used to determine the number of clusters by picking the higher average, in the case of comparing two candidate clusterings.

**Figure 4.**
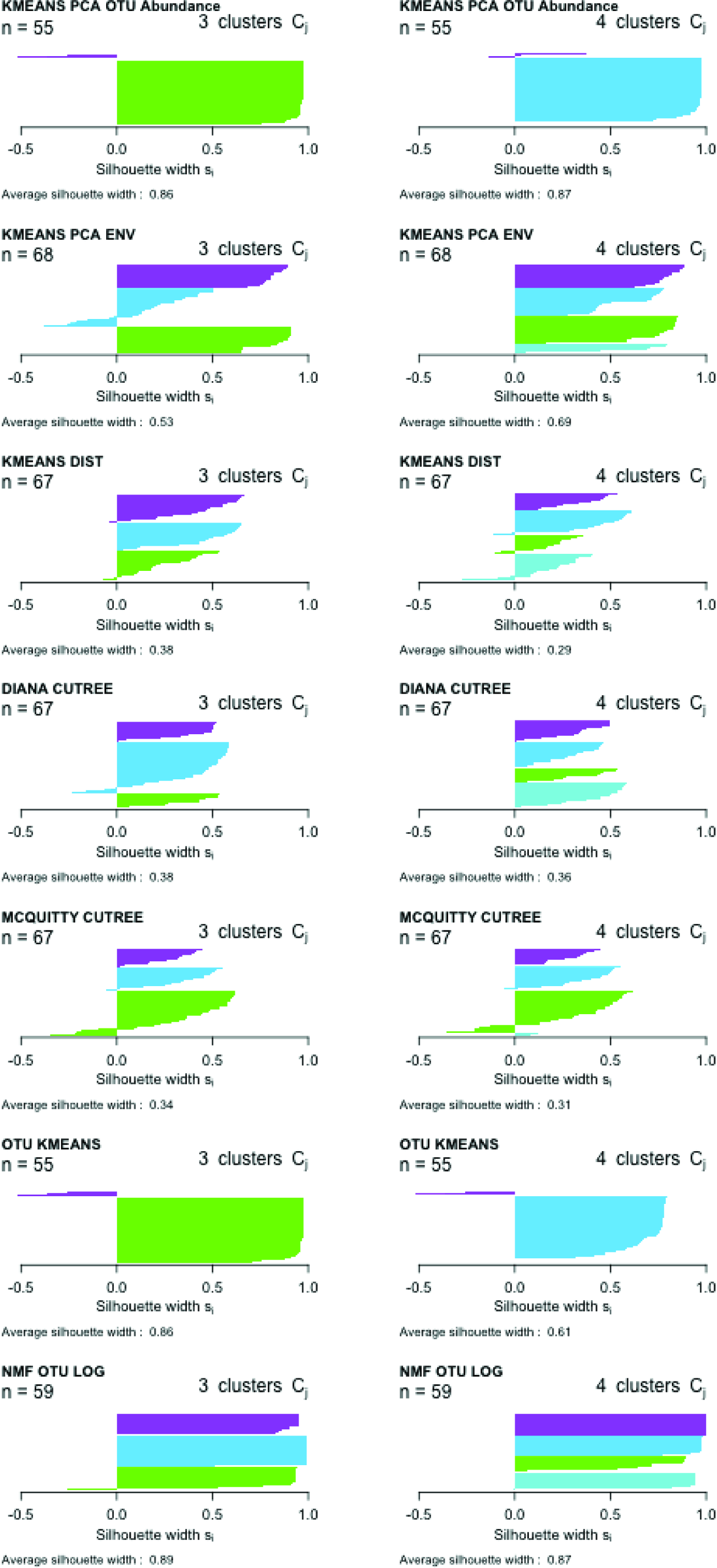
*Left side has 3 clusters, while right side utilizes 4 clusters. Top to Bottom: PCA determined clusters on OTU abundance, PCA determined environmental clusters, kmeans on mash distances, diana on mash distances, kmeans on OTU data (euclidean distance), NMF on OTU data, K=3 and k=4 were found to be viable, on the determination that an Average Silhouette Width above 0.5 was acceptable. A score above 0.25 may indicate structure* (Rousseeuw, 1987)*. The environmentally driven PCA produced viable clusters at both K=3, and K=4, Kmeans on sequence similarity at K=3 weakly indicated structure, Diana clustering weakly indicated structure at K=3 and K=4, and NMF on OTUs produced structured clusters at both K=3 and K=4. All Silhouette Widths show that clustering at 3 or 4 maybe viable, with the exception of K-means on OTUs, in which most samples clustered into a single, large group. Not all samples were viable in each method, “n=” indicates number of samples utilized in each method.*

Because Silhouette Width Average for different clustering methods fell at the best values at either K=4, or at K=3, new clusters were generated at both. At K=3, clusterings were generated by Random Assignment, Nonnegative Matrix Factorization based on abundance information, as well as log transformed OTU abundance, the three clusters with least within cluster distances from both the Diana, and from Mcquitty-WPGMA hierarchical clustering. Clusters were also made from Sample PH and from a North, South, or Central location. Since environmental variable mixing was previously reported to be the driver of beta diversity at k=3 (Crits-Christoph et al. 2013), we used environmental variable mixing to also generate clusterings. K=4 clusterings were generated with Random Assignment, Non-negative Matrix Factorization based on abundance information, as well as log transformed abundance information, the four clusters with least within cluster distances from both the Diana, and from Mcquitty-WPGMA hierarchical clustering, as well as from PH. Cluster to Cluster correlations show that Mcquitty-WPGMA is more similar to environmental clusterings; however, all non-random clusterings are more similar to each other than to randomly generated clusterings, indicating all detect some elements of community structure present in the data. Although this analysis has indicated that there was an ecological correlation to computationally derived clusters, it has not shown which factors, or how those factors affect clustering. Further, skewed species abundances with a few dominant species could make it more difficult to sample rare species at modest sequencing depth; however, because Mash estimates the similarity between two sets, slight stochastic differences in observed abundances should not significantly affect the results relative to traditional OTU approaches that are also subject to

**Figure 5.**
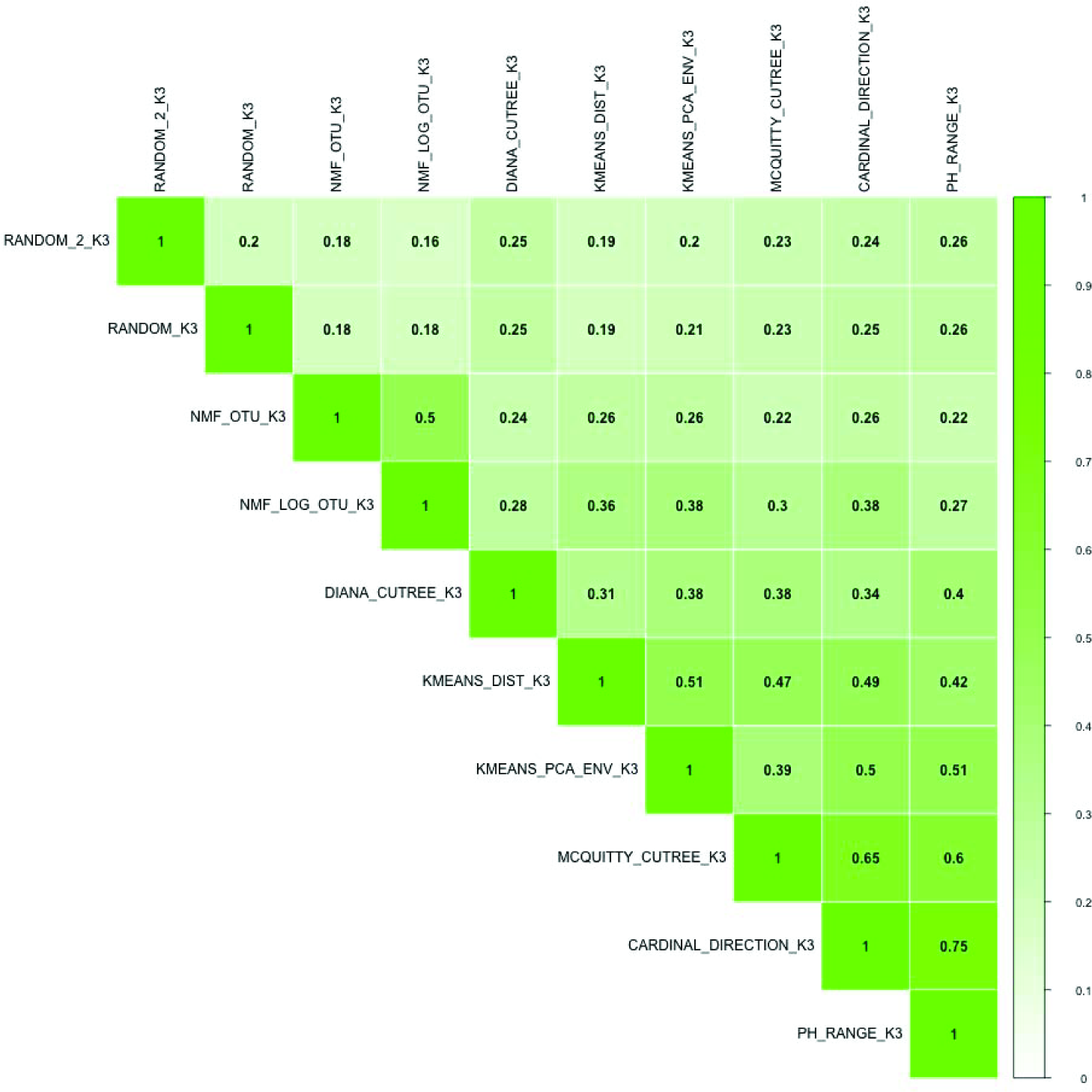
*represents cluster similarities between each cluster, cluster similarity jaccard algorithm was used, k=3 clusterings are shown.*

**Figure 6.**
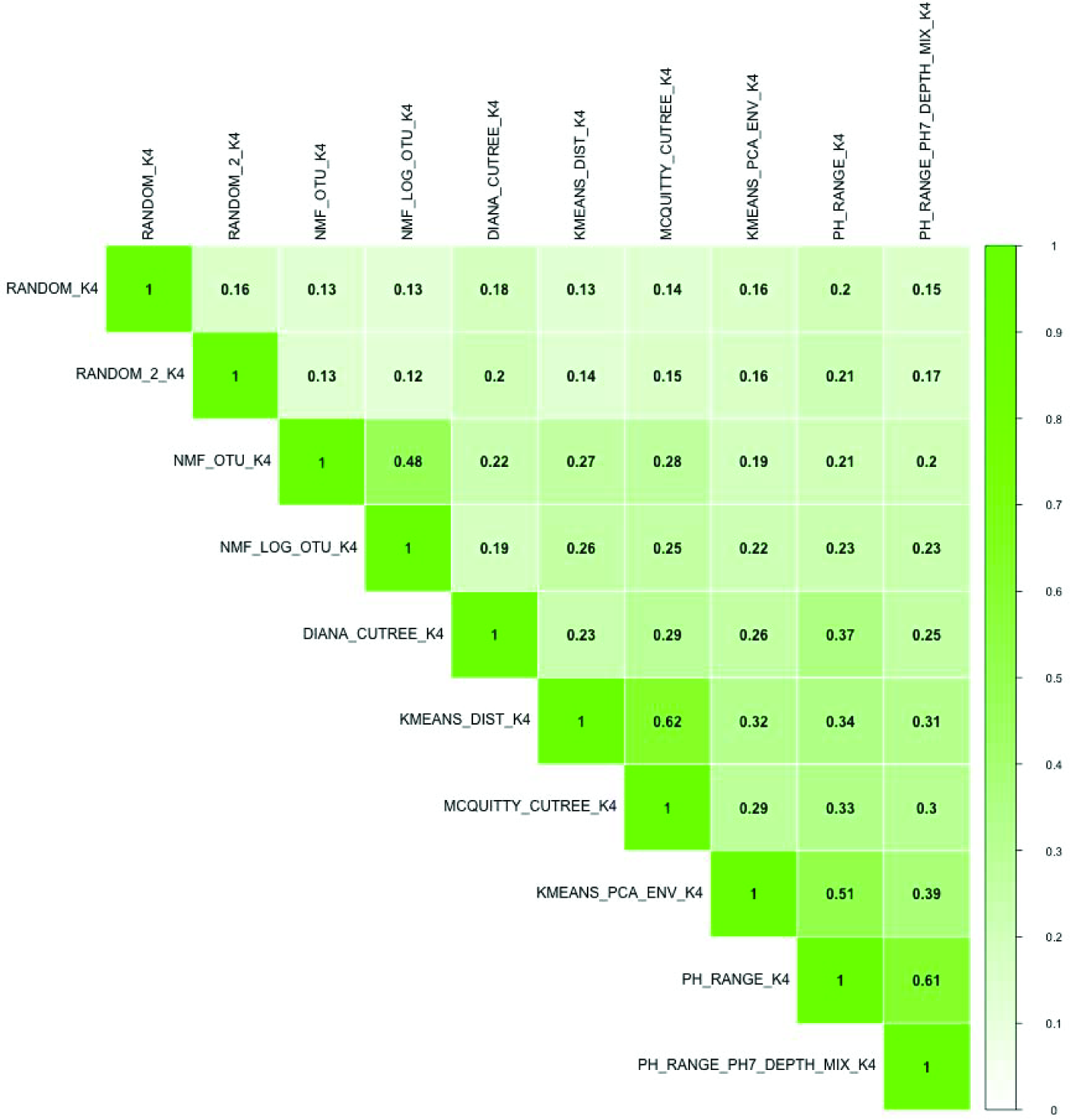
*represents cluster similarities between each cluster, cluster similarity jaccard algorithm was used, k=4 clusterings are shown.*

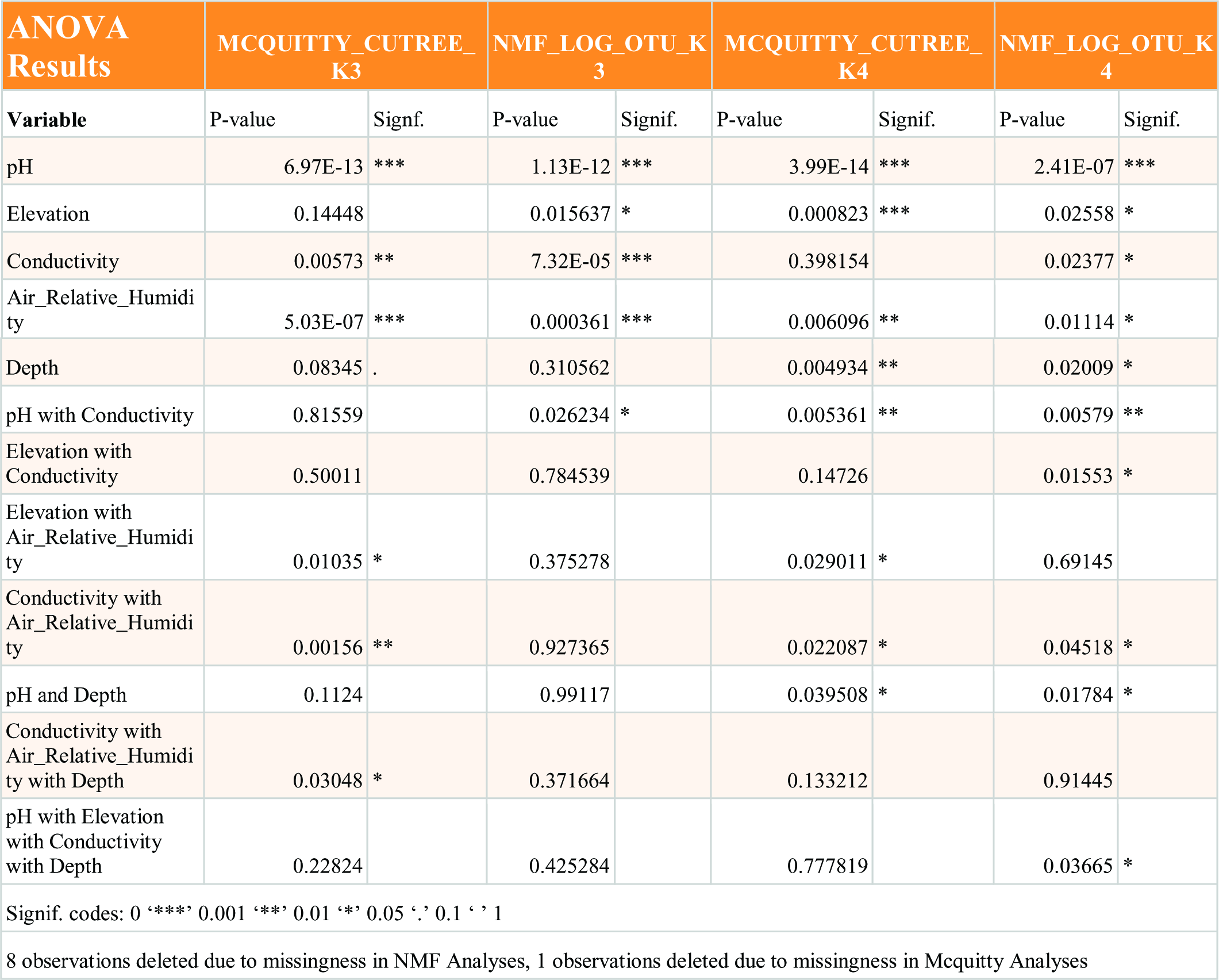

The clusterings were modeled by ANOVA, and after calculating a log likelihood test, we found that for both K=3 and K=4 Mcquttity hierarchical clustering, followed by NMF on OTUs, were the most significant and therefore best corresponded to the environmental data. For McQuitty hierarchal clustering, PH and Elevation were found to have the most significance, however, since the elevation was the same for all of the samples of any given sampling site, and since elevation is highly correlated with sampling location there may be some other latent variable that is being indirectly measured, also highly correlated with sampling location. For NMF on log abundance PH, Conductivity, and Relative humidity of the air were found to be most significant; however, because relative humidity of each sampling site was the same, it is unknown whether relative humidity of the air was the contributing factor or some other, unknown variable, that also differed from site to site was a factor.

McQuitty clustering has a .65 similarity with the Cardinal Direction, and similarly high similarities with other environmentally determined groupings. We also see that both McQuitty and NMF have high p-values with some environmental variables in anova, with Ph being particularly significant in both McQuitty and NMF, and to a lesser extent Relative Air Humidity being significant as well, and that the sample similarities within these groupings are high. This shows that OTU based methods and distance-based methods produce similar results, if driven by slightly different environmental variables, and is getting at the underlying structure of the local communities.

As per the clustering Silhouette Widths, some of the methods, Diana and NMF, work better at four clusters, while McQuitty and K-means did better at three. The most explicative results, as per ANOVA, NMF on log OTU abundance and McQuitty slightly disagree on which environmental variables have the most importance, but PH and Relative Air Humidity can be seen across all four ANOVAs.

## Discussion

How communities are determined, or even how they are considered, is up for debate. Are communities composed of nearly homogenous samples or are they composed of a mix of different kinds becomes an important question that drives experimental design (Holyoak et al. 2005). If the expectation is that samples should be nearly homogenous, determining communities algorithmically via sketch-based clustering is possible. In this study, however, we observed that samples’ clusters derived from MASH had low cluster to cluster similarities relative to the derived clusters using OTUs/Mothur, even with quality trimmed data supplied to MASH. One likely explanation is MASH-driven clustering is being affected by sequences not deemed as OTU sequences either because of low coverage, contamination, insufficient length, or some combination therein in the read data published previously. Even so, ANOVA analysis on the derived clusters showed that some measured environmental variables were consistently significant while others depended on the clustering method used, with MASH and NMF having the high associations. When considering NMF as an alternative this may make some sense given that it is a factorization method focused separating two clusters, while traditional PCA would focuse on variables with the most divergence between samples. As such NMF factors can be intuitively understood and can allow for an overlap in basis components (Gaujoux & Seoighe, 2010) making it a viable choice in determining communities.

Finally, we believe that abundance derived *de novo* clusters are useful specifically because they group samples without prior knowledge of geospatial or other overriding factors. This is powerful to assess associations between species abundance, communities, and environmental variables (inc. geospatial) without requiring a more complex statistical model.

## Concluding Remarks

**Figure 9.**
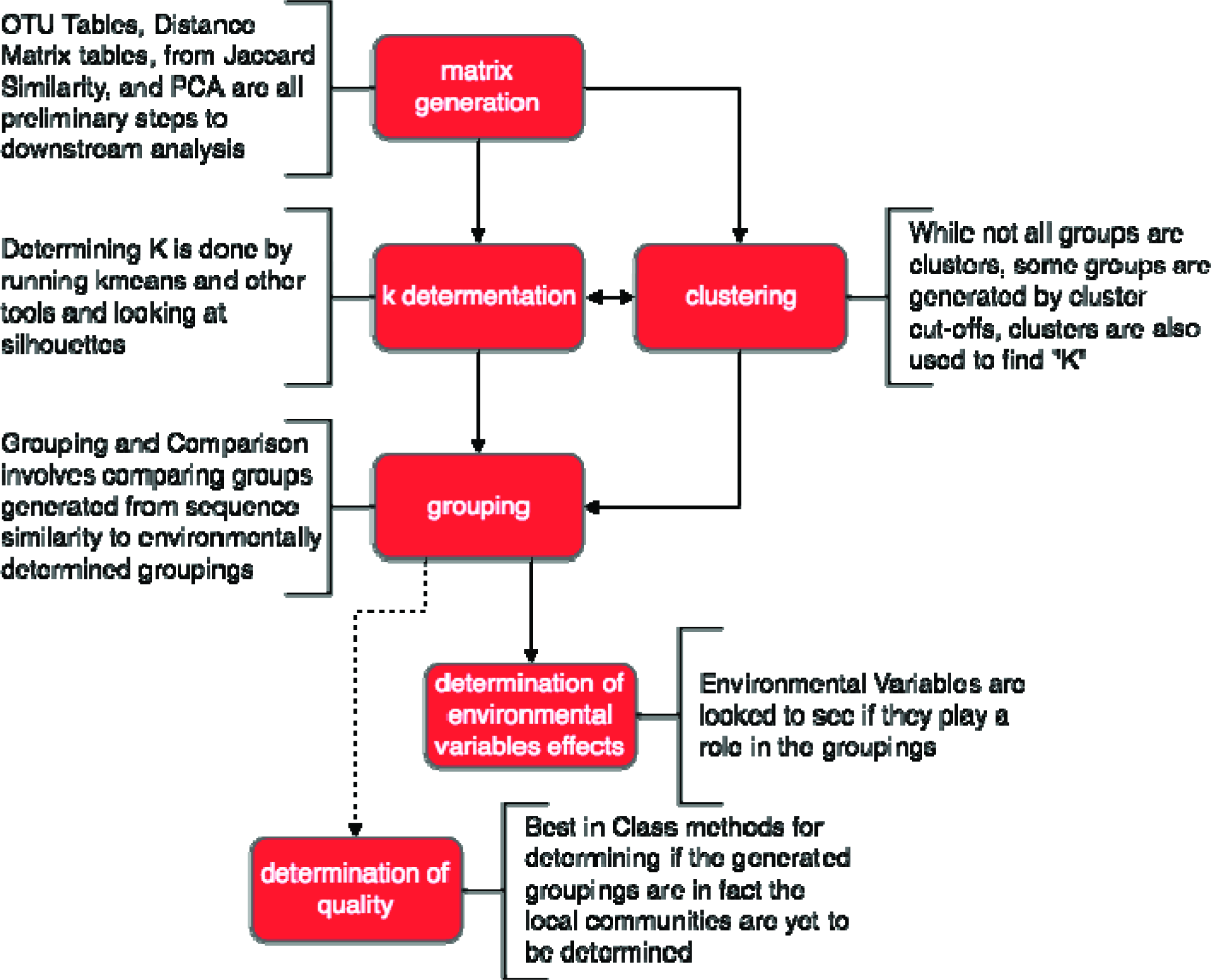
Workflow for Determining, Describing, and Validating Atacama data.

As a general workflow (figure 9), after sample collection either OTUs abundances are generated, or sample to sample distances are calculated by comparing their contained trimmed sequences. In the case of the sample to sample distances a distance matrix is generated that can be clustered though hierarchical or other means, and in the case of OTU abundances NMF or K-means is better suited. We calculated pairwise distance on both shared ple to to sequences and on OTUs, and then clustered OTUs and shared sequences via K-means, and for shared sequences diana clustering was also utilized, and for NMF was also utilized for OTU abundance. The groupings can then be checked for the influence of independent variables, through a statistical model, in this case anova, which was run on clusters, the ‘anova’ function from the R ‘stats’ package was used, and LogLik from the R ‘stats’ package was used to compare Log-Liklihoods. Clusters were compared to each other using both RAND and Jaccard similarity cluster evaluation methods, as well as a wilcox test (Hollander et al. 2013); (Bauer, 1972).

The Atacama data used here is from SRA:SRA091062, Bioproject ID: PRJNA208226, which was thought of as three clusters of data, aligning with sampling site: North, Central, and South. Atacama was chosen for its previous environmental analysis, geographically distinct sampling sites, and curated metadata.

Mothur was used to process Raw files for OTU analysis as per non-shhh (Quince et al. 2009) 454 SOP: https://www.mothur.org/wiki/454_SOP. for sequence similarity distance Mothur was used to filter samples based quality scores, as per the shhh and trimming portion of the mothur 454 SOP. Initial NMF analysis (figure 10) was done with “sake” (https://github.com/naikai/sake), which was originally created to analyze gene expression data, was here utilized to look at OTU abundance data, at k=3 both log transformed and non-log transformed data was utilized, the nsNMF NMF algorithm NMF algorithm was used and the the NMF tool was run at 350 runs, at k=4 only log transformed data was run, with the nsNMF (Pascual-Montano et al. 2006), NMF algorithm at 350 runs. nsNMF was chosen for its design to deal with perceived sparseness in the data. The R ‘cluster_similarity’ function from the ‘clusteval’ package was used for Jaccard and RAND similarities, while ‘wilcox.test’ function from the R ‘stats’ package was used for the wilcox test. wpgma was chosen for Agglomerative clustering because clusters were expected to be of unequal size, as unweighted hierarchical methods can become distorted when large and small groups are compared, and a clear contrast to centroid clustering, as like k-means, was desired. The R ‘hclust’ function was used from the ‘stats’ package was used for agglomerative clustering. Diana, from the R ‘cluster’ package was used for divisive hierarchical clustering, in agglomerative hierarchical clustering samples are combined until all samples are in the same cluster, whereas in divisive hierarchical clustering all samples start in the same cluster and then are partitioned into daughter clusters.. And further analysis and figure analysis was done with the caret (https://cran.rproject.org/package=caret), clusteval (https://cran.r-project.org/package=clusteval), cluster (https://CRAN.Rproject.org/package=cluster), corrplot (https://CRAN.R-project.org/package=corrplot), d3heatmap (https://CRAN.R-project.org/package=d3heatmap), fpc (https://CRAN.R-project.org/package=fpc), gplots (https://CRAN.R-project.org/package=gplots), and NMF (https://CRAN.R-project.org/package=NMF) R packages.

Supplemental files (https://github.com/status-five/Methods-in-Description-and-Validation-of-Local-Metagenetic-Microbial-Communities/releases/tag/v1.0) (Molik, 2018)

